# Dissociable Roles of FEF Neurons in Initiating Saccades

**DOI:** 10.1101/2022.02.01.478650

**Authors:** Mohammad Shams-Ahmar, Peter Thier, Yaser Merrikhi

**Affiliations:** Cognitive Neurology Laboratory, Hertie Institute for Clinical Brain Research, University of Tübingen, 72076 Tübingen, Germany; Department of Physiology, McGill University, Montréal, Canada

**Author notes:** Correspondence: Mohammad Shams-Ahmar.

## Abstract

The frontal eye field (FEF) plays a key role in initiating saccades. To explain how single neurons in this region may enable the initiation of saccades, Jeffery Schall proposed the “variable rate model” (Hanes and Schall, 1996) which assumes that the discharge of a neuron must reach a fixed activation threshold for a saccade to be evoked. However, the validity of this model has been questioned as results from later work testing saccades in different behavioral contexts seemed to require the assumption of variable thresholds. Moreover, even when sticking to the original behavioral context, not all types of frontal eye field neurons seemed to behave according to a variable rate model. In an attempt to reconsider the viability of the variable rate model avoiding limitations of previous research, we studied single neurons recorded from the FEF of two rhesus monkeys while they performed a memory-guided saccade task. We evaluated the degree to which each type of FEF neurons complied with the variable rate model by quantifying how precisely their discharge predicted an imminent saccade based on their immediate presaccadic activity. In addition, we asked whether there might be a hierarchical relation between visuomotor and motor neurons with only the latter responsible for the release of saccades and therefore satisfying the assumption of the variable rate model. We show that decoders trained on single motor neurons’ presaccadic activity performed better than decoders trained on visuomotor neurons in predicting an imminent saccade, in line with the assumption of a top position in a putative processing hierarchy. While this position might have suggested significant resilience to perturbation of the visual input to the FEF, we found quite the opposite that motor neurons but not visuomotor neurons were susceptible to perturbation. As a matter of fact, they lost the ability to predict the initiation of a saccade based on their immediate pressacadic discharge rate. Our results support the notion that motor neurons and visuomotor neurons have dissociable functions in information processing for saccades with motor neurons more closely related to saccade initiation, in line with the tenets of the variable rate model. Yet, they also clearly indicate that even these supposedly top layer neurons exhibit an unexpected degree of dependence on afferent input questions the viability of the assumption of a simple processing hierarchy.

## Introduction

A causal role of the frontal eye field (FEF) in initiating and overall in generating saccades has been demonstrated in two ways: Electrical microstimulation of this region evokes saccades (Bruce et al., 1985; Opris et al., 2001; Robinson and Fuchs, 1969) and its pharmacological inactivation impairs the generation of saccades (Dash et al., 2018; Sommer and Tehovnik, 1997). To explain how single neurons in this region could contribute to saccade initiation, (Hanes and Schall, 1996) suggested a “variable rate” model. According to this model, different rates of accumulation of neural activities at the single-cell level would result in different initiation times by reaching a fixed threshold at different times. The model implies that shortly before saccade onset, neurons with saccade-related activity will discharge at the same fixed level, independent of the time of saccade onset relative to a go-cue signal.

To test whether neurons reach a fixed level of activity before saccade onset independent of saccade time, several previous studies resorted to a correlation analysis of the relationship between saccadic reaction time and discharge rate immediately before the saccade (Fecteau and Munoz, 2007; Hanes and Schall, 1996; Hauser et al., 2018; Jantz et al., 2013; Paré and Hanes, 2003), guided by the expectation of independence between the reaction time of saccades and the presaccadic activity level resulting in a correlation indistinguishable from zero. However, the viability of this analytical approach depends on three important prerequisites: first, the number of samples in the correlation analysis must be large enough to ensure sufficient statistical power to detect deviations from the null hypothesis — the absence of a correlation; second, it needs to be shown that activation thresholds are reliable enough to allow a precise prediction of an imminent saccade; and third, it must be shown that the activation level has not already been reached prior to the temporal window that comprises information indeed responsible for saccade initiation.

A problem for the variable rate model as proposed originally is that later work suggested that specific cognitive demands seem to call for a change of the putative saccade initiation threshold thought to be given by the level of presaccadic activity of FEF neurons. For instance, it has been reported that the presaccadic activity needed to release a saccade was higher for pro-saccades than for anti-saccades (Everling and Munoz, 2000; Jantz et al., 2013); also, presaccadic activity was higher when the instruction called for a fast response rather than an accurate one (Reppert et al., 2018). Moreover, different classes of FEF neurons seem to be differently susceptible to these varying demands. In general, motor neurons are more compatible with the assumptions of the variable rate model than visuomotor neurons (Jantz et al., 2013). This suggests the possibility that motor neurons might be the primary mediators of saccade initiation, while at the same time, raising the question of what the contribution of visuomotor neurons might be.

In this study, we tested the viability of the variable rate model separately for FEF motor and visuomotor neurons recorded from two rhesus macaques during a memory-guided saccade task. First, we found that motor neurons support this model much better than visuomotor neurons when deploying correlation techniques. Second, to avoid insufficiencies of the correlation approach mentioned above, we ran a complementary decoding analysis in an attempt to challenge the variable rate model. The idea was that if the model holds, a decoder that compares activity very close to saccade onset with activity produced earlier should be able to predict the occurrence of a saccade. In fact, the result of the decoding analysis supported the conclusions of the correlation analysis. Finally, assuming that motor neurons — and not visuomotor neurons — are instrumental in initiating a saccade, the code extracted by the classifier should be much more resilient to perturbations to the FEF. This expectation was tested by microstimulating V4, one of the major inputs to the FEF (Schall et al., 1995; Stanton et al., 1995). However, at odds with our expectation, we found that the saccade code of the motor neurons was highly sensitive to microstimulation. We conclude that motor neurons’ contribution to saccade initiation can be sufficiently explained by the variable rate model. However, the fact that motor neurons, supposedly representing an output layer, exhibited an unexpected degree of dependence on afferent input is at odds with the concept of a simple processing hierarchy.

## Methods

### Data acquisition

The data was recorded from two male rhesus monkeys (Macaca mulatta). All experimental procedures were in accordance with National Institutes of Health Guide for the Care and Use of Laboratory Animals, the Society for Neuroscience Guidelines and Policies, and Stanford University Animal Care and Use Committee. The surgical and the data acquisition procedures were described in detail elsewhere (Noudoost et al., 2021). In brief, the electrophysiological data were recorded extracellularly from the frontal eye field (FEF) using a multi-site, linear electrode (v-probe Plexon) and the receptive fields (RF) were mapped and later confirmed by electrical microstimulation evoked saccades (≤ 50 μA, 250 Hz; 0.25 ms duration) made toward the center of the mapped RF.

After having calibrated the eye position records provided by a video eye tracker (500 Hz), the monkeys were required to perform a memory-guided saccade task (MGS): they were asked to fixate a round one degree fixation dot for one second. Then a cue (squared shape, 1.35 deg) appeared either at the center of the RF or at the mirrored position in the opposite hemifield outside the receptive field for one second. The cue disappeared and after another one second the fixation dot went off, providing a go signal for the monkey to make a saccade within 500 ms toward the remembered location of the cue, with an acceptable spatial error window of four degrees.

During half of the trials, electrical microstimulation was delivered to area V4 at one of the four potential time points during the task. The electrical microstimulation was a single biphasic current pulse (600–1000 μA; 0.25 ms duration, positive phase first).

### Data analysis

The presence of visual activity was tested by demonstrating a significant difference (p < 0.01) between the mean discharge rate in the last 300 ms of the baseline fixation period before cue onset and an interval of 300 ms starting 50 ms after cue onset (visual period). The presence of saccade-related activity was tested by demonstrating a significant elevation of the discharge rate in a perisaccadic window of 100 ms window duration, centered on saccade onset relative to the mean discharge rate in a 100 ms window starting 50 ms after fixation dot disappearance (the go signal). We limited the scope of our analyses to neurons with at least a modulation during the perisaccadic period in trials, in which the target appeared in a neuron’s response field. Neurons with a modulation during the perisaccadic period only were considered as motor neurons, and neurons with a modulation during both saccade and visual periods were considered as visuomotor neurons.

Regressions between spiking activity at varying times within a window from −70 ms to saccade onset and saccadic reaction time were based on a nonparametric approach (Kendall’s tau), using a translating window of 10 ms starting at a given time point (forward averaging).

In order to address the question if at a particular time before the saccade the discharge would be able to predict an ensuing saccade, a decoding analysis was employed. We defined two classes of presaccadic bins: a fixation bin and a saccade bin. The fixation bin at a given time point was the spike count over a 10 ms window, randomly selected from a larger window of 100 to 10 ms before that time point. The saccade bin was the spike count over a 10 ms window starting from the time point and ending 10 ms later. For each neuron and pair of bins, a linear classifier (LDA) was trained to predict the occurrence of a saccade. To this end half of the trials, randomly selected from all trials of a given neuron were used for the training and the resulting classifier was then tested with the remaining half. This procedure was repeated 25 times and the average prediction performance on test trials was taken as a measure of the final performance of the neuron.

## Results

A total number of 123 FEF neurons with presaccadic activity were collected from two monkeys (Fig. 1A) performing a memory-guided saccade task (Fig. 1B). We identified 59 (48%) motor neurons and 64 (52%) visuomotor neurons (see Methods). Figure 1C demonstrates the average firing rates of two exemplary neurons. The discharge pattern of the sample motor neuron (top) shows a strong modulation in the saccadic period, peaking almost at the time of saccade onset. The sample visuomotor neuron exhibited bouts of activity at the onset and offset of the target cue appearing in its receptive field in addition to a broader activity peak around the time of the saccade.

**Figure 1.**
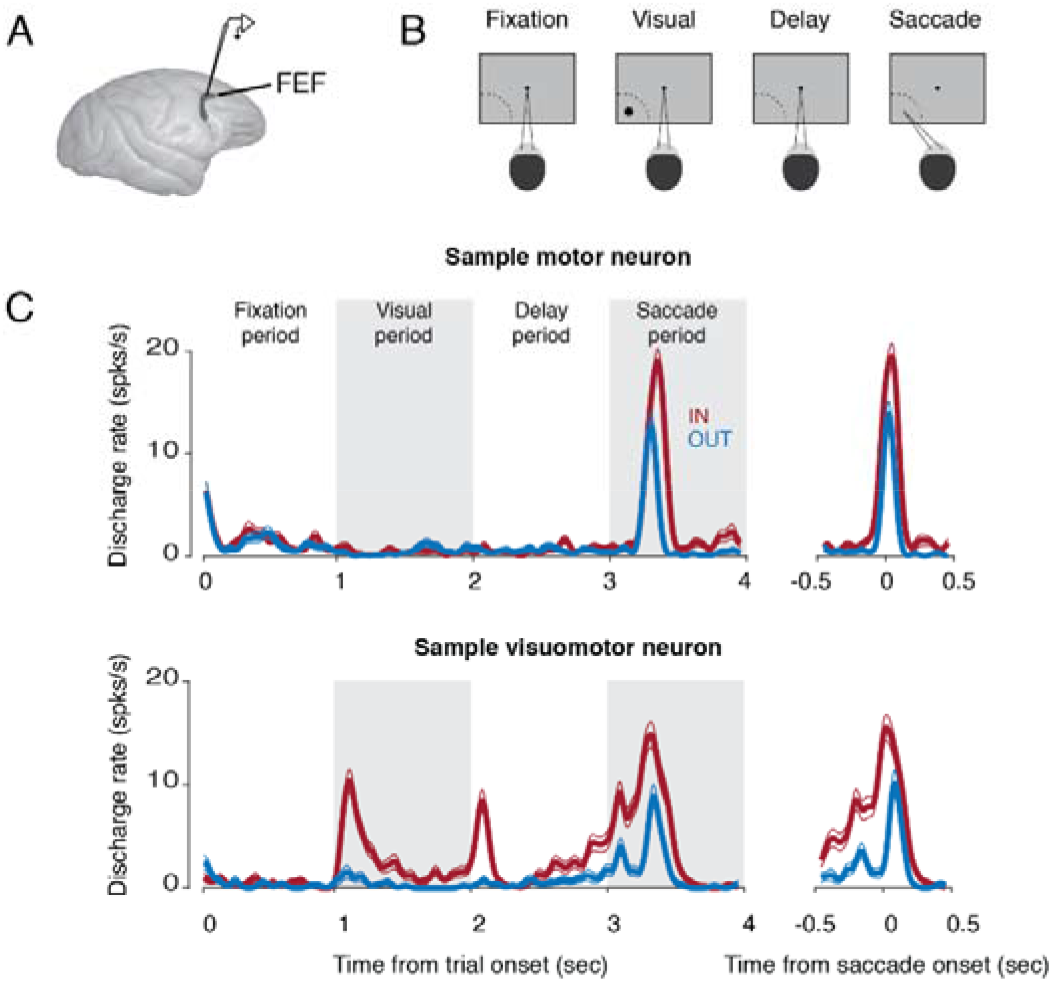
Behavioral paradigm and exemplary single neurons. A) Recording site. B) An example of the IN condition of the memory guided saccade task. The monkey fixated a spot in the center of the screen for one second after which a target cue appeared inside the motor field (dashed curved line) for another one second. The monkey kept fixating the central dot until its disappearance signaled the monkey to make a saccade toward the memorized location of the target cue. C) Average discharge pattern of an exemplary motor neuron (top) and visuomotor neuron (bottom) aligned to fixation onset (left) and aligned to saccade onset (right). Thick colored lines indicate the average firing rate across trials and the thin colored lines indicate ± s.e.m (red: IN condition; blue: OUT condition). Shaded gray regions are for better visual dissociation of the four task periods.

Figure 2A gives a schematic illustration of the relation between activation threshold and reaction time as predicted by the variable rate model. According to this model, after a go-cue — provided by the disappearance of the fixation dot in our experiment — the discharge rate of an FEF neuron starts to build up until it reaches a fixed threshold (horizontal dashed line). Faster build-up rates entail that the threshold will be reached earlier and a saccade will be elicited that has a shorter reaction time. As the model assumes a constant threshold, saccade latency must be independent of discharge rate at the time a saccade is released. Hence, the correlation between saccadic reaction time and the discharge rate right before the saccade must be zero. Figure 2B illustrates an alternative scenario, in which variable build-up rates do not terminate at the same threshold. In this particular example, the threshold decreases over time (oblique dashed line). This would entail that slower discharge build-up rates would terminate at lower thresholds, ultimately resulting in a negative correlation between the immediate presaccadic discharge rate and the saccadic reaction time.

**Figure 2.**
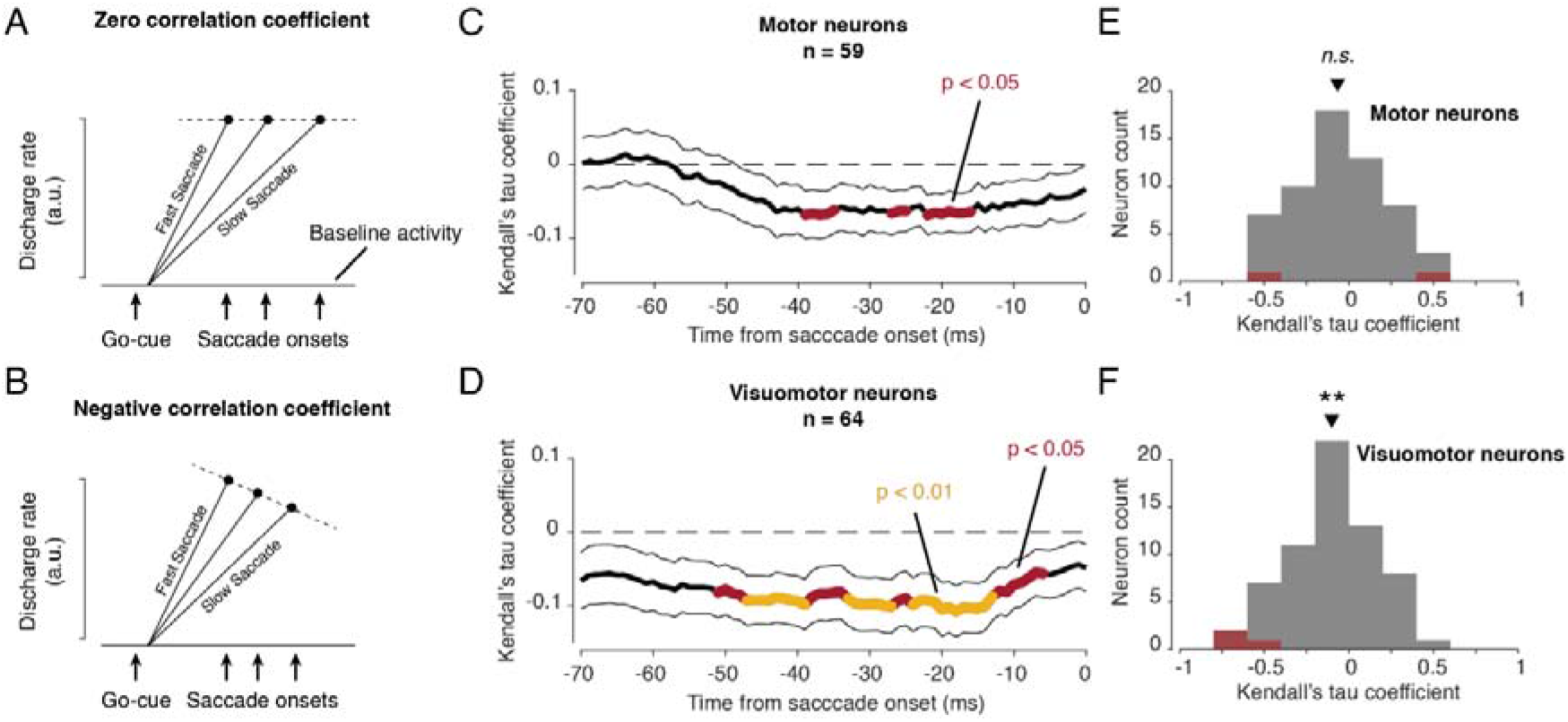
Relationship between presaccadic activities and reaction times. A) Schematic illustration of the variable rate model. After the go-cue signal, the activity of an FEF neuron builds up to a fixed threshold (activation threshold) at which the saccade is released. A faster build-up of the discharge results in faster saccades and vice versa. Here the correlation between the discharge rate at threshold (i.e., at a fixed time shortly before saccade onset) and reaction time is zero as indicated by the horizontal dashed line B) An alternative model. Here the threshold is assumed to change with time. The hypothetical threshold profile would entail that slower discharge build up rates terminate at lower thresholds, hence resulting in a negative correlation between immediate presaccadic discharge and reaction times as indicated by the oblique dashed line. C-D) Correlation time course (mean ± s.e.m) based on a 10 ms moving window for motor neurons (C) and visuomotor neurons (D). Colored lines indicate time-points at which the average correlation of the population of neurons was significantly different from zero (red: p < 0.05; orange: p < 0.01). E-F) Kendall’s tau distributions based on activity in the activation window as proxy of the activation threshold of motor neurons (E) and visuomotor neurons (F). Neurons with significant correlation coefficients in the activation period (p < 0.05) are shown in red. *n.s*. indicates p ≥ 0.05, ** indicates p < 0.01.

In order to investigate if the assumption of a fixed activation threshold, independent of the reaction time, posited by the variable rate model, holds, we first looked at the correlation of discharge rate and saccadic reaction time. Since it is not clear which time window preceding saccades represents the threshold, relevant for the generation of saccades, we calculated the correlation for 10 ms bins starting 70 ms before saccade onset until the time of saccade. Figure 2C-D show the average correlation between reaction times and activation thresholds across neurons.

In accordance with the assumption of a variable rate model with fixed threshold, the average correlation coefficient of motor neurons rarely deviated significantly from zero (Fig. 2C). Conversely, visuomotor neurons consistently deviated from zero in a large window starting at about 50 ms until shortly before saccade onset (Fig. 2D), contrary to the tenets of a variable rate model. A similar conclusion was suggested by the distribution of the correlation coefficients of neurons for a window of −50 ms to −40 ms before saccade onset, a window that is presumably relevant for saccade initiation (from now on referred to as “activation window”; Bruce et al., 1985; Opris et al., 2001; Robinson and Fuchs, 1969). Whereas the distribution for motor neurons (Fig. 2E) did not deviate significantly from zero, the visuomotor neurons (Fig. 2F) exhibited a significant deviation (p < 0.01) because of a dominance of neurons with negative correlation coefficients, no matter if significant at the single neuron level or not.

These results might be taken as support of the conclusion that FEF motor neurons act according to the tenets of the variable rate model with fixed threshold. Yet, as discussed earlier, the correlation approach has limitations and might lead to false positive conclusions. Therefore, we resorted to a decoder-based approach described in the Methods section in order to clarify if discharge in a window (the “saccade class bin”) with a specific temporal distance to the later saccade is able to predict it. As illustrated in Fig. 3A this would be the case if the discharge in the saccade bin could be reliably separated from the discharge level characterizing activity in the preceding fixation bin. However, in case of too large variability also the decoder would fail (Fig. 3B). However, the decoder approach will in any case be able to avoid the problem of false positives due to too small sample sizes.

**Figure 3.**
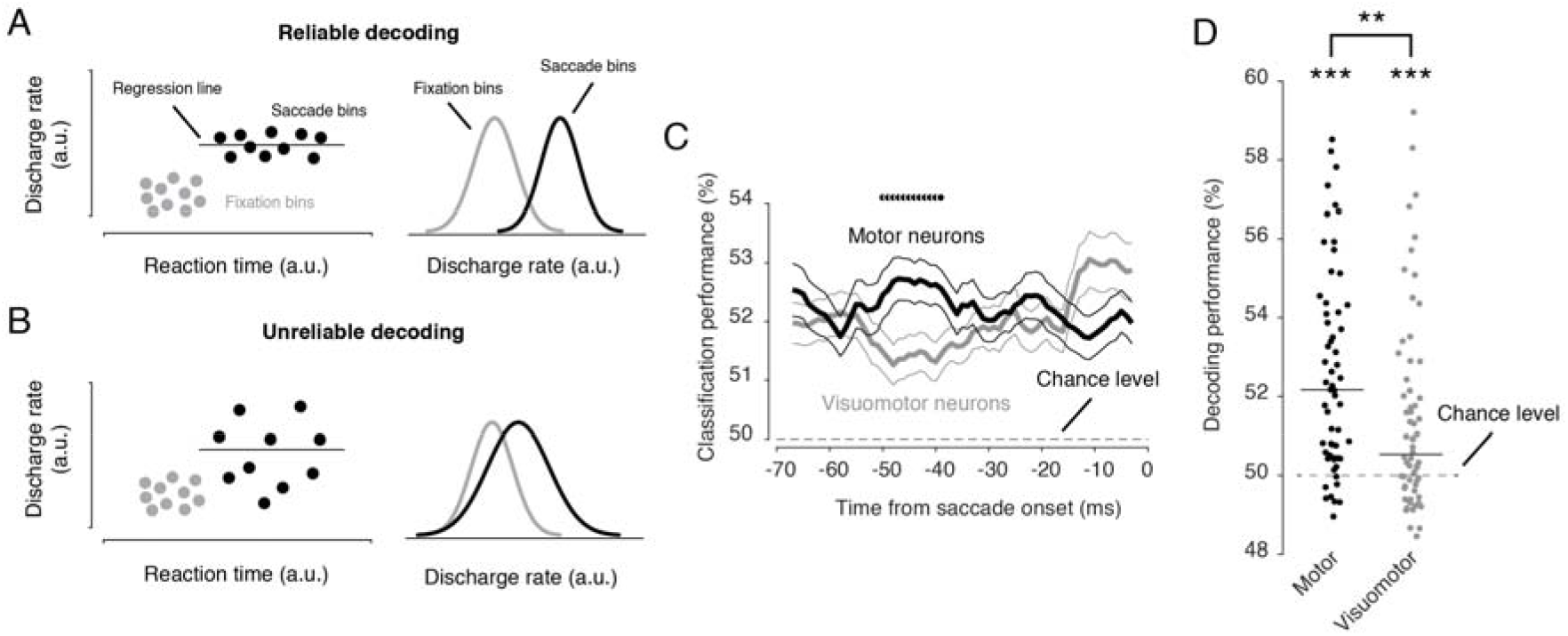
Reliability of the presaccadic code. A-B) Two illustrative examples of a relation between reaction time and activation threshold in an FEF neuron. Gray circles represent fixation bins as a proxy for the discharge associated with fixating eyes. Black circles represent saccade bins as proxy for the discharge responsible for saccade initiation. Both examples (A and B) demonstrate a zero correlation, but in (A) a decoder would be able to reliably decode the time of saccade based on the activity levels, but in (B), a decoder would perform poorly, questioning the interpretation of a zero correlation. The horizontal gray lines are the regression line with a slope of zero. C) Time course of the actual performance of a decoder trained on half of the trials and tested on the other half for each FEF neuron. Black circles indicate time points with significant difference (p<0.05) between the performance of motor neurons and visuomotor neurons. The results are shown separately for motor neurons (black) and visuomotor neurons (gray). D) The average performance of each neuron over a window of −50 to −40 from saccade onset (the “activation window”) is shown separately for each FEF neuron type. Each dot represents one neuron. Horizontal solid lines show medians. ** indicates p < 0.01, *** p < 0.001.

We determined the time course of the reliability of the saccade prediction within a window from 70 ms before to the time of saccade onset as described in detail the Methods sections by comparing activity in two subsequent bins whose position was shifted with this window. We argued that if the activity in the leading saccade class bin were a reliable marker of saccade initiation, then the decoder, comparing it with activity in the preceding fixation class bin should be able to reliably predict the occurrence of a saccade.

Figure 3C shows the resulting time course of the decoder performance analysis, separately for motor and visuomotor neurons. About 50 ms before saccade onset, i.e., within the “activation window” the performance for motor neurons started to increase while the visuomotor neurons decreased at the same time, however still staying above chance level. This difference in the performance trajectories for the two types of neurons was significant for more than ten consecutive time points. Also, after averaging the predictions over the activation window both types performed above chance (Motor: mean = 52.2%, p < 0.001; Visuomotor: mean = 50.5%, p < 0.001). However, the information provided by motor was clearly more reliable than the one for visuomotor neurons (p < 0.01).

Finally, we asked whether motor neurons and visuomotor neurons were differently susceptible to disruptions. To this end, while recording from FEF during the memory-guided saccade task, in half of the trials (selected randomly), we delivered electrical microstimulation at sites in area V4 representing congruent regions of the motor field (Fig.4A). Stimulation was delivered at four different time points as specified in fig.4B. We used the same decoders trained on no-stimulation trials, described in the previous section, and tested them this time on the stimulation trials. If the motor code did not change as a result of microstimulation, the performance of the decoders should not drop, compared to their performance for no-stimulation trials.

**Figure 4.**
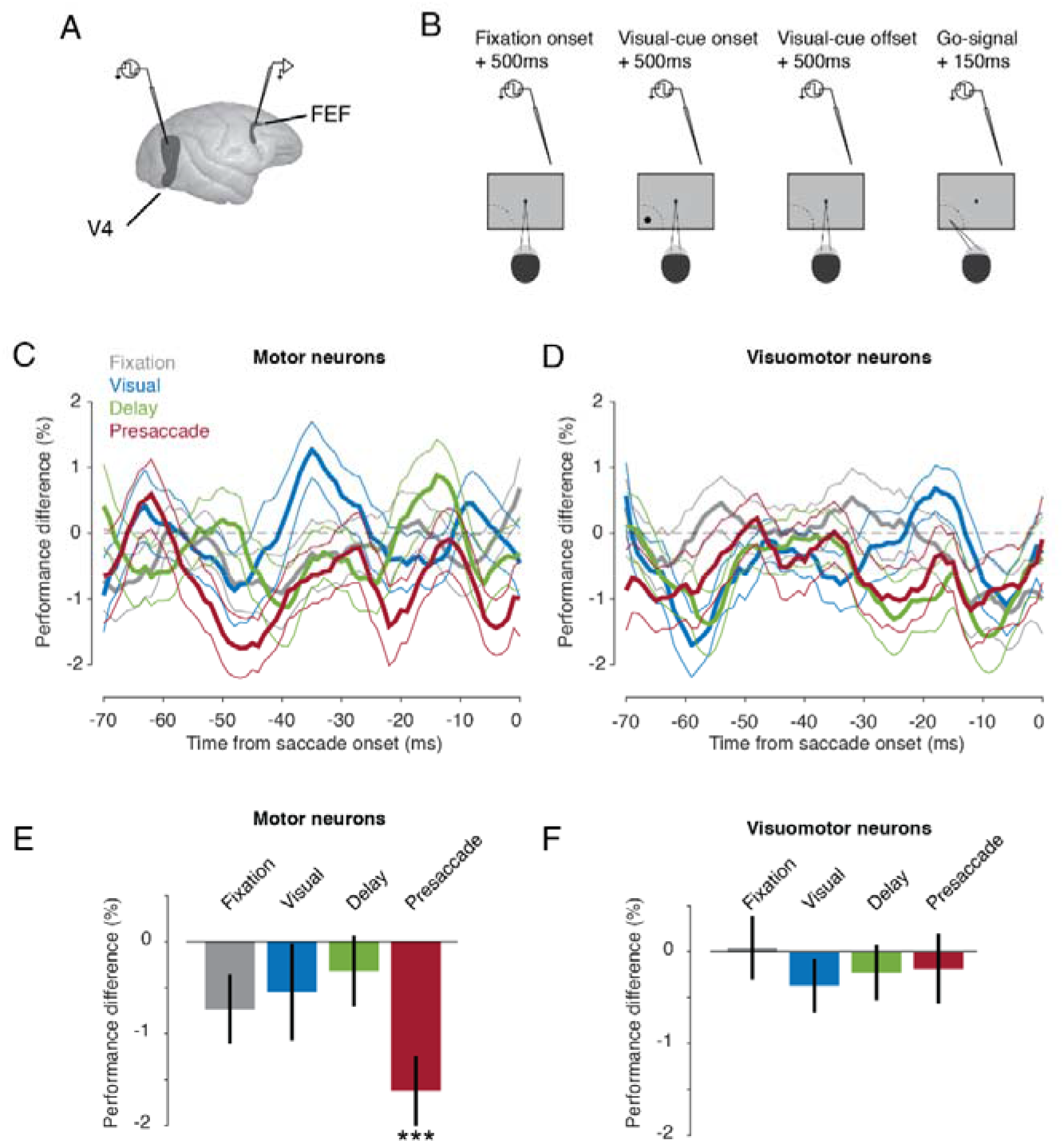
Effect of microstimulation on the saccade code. A) Microstimulation and recording sites. B) While recording from FEF, in half of the trials, electrical microstimulation was delivered to area V4 on the same hemisphere. The microstimulation was delivered 500 ms after fixation onset, 500 ms after visual-cue onset, 500 ms after visual-cue offset, or 150 ms after the go-signal. C-D) The average time course of the difference of decoding performance for stimulation and no-stimulation trials for motor neurons (C) and visuomotor neurons (D). At each time point, a decoder was trained based on the sum of a 10 ms window (forward moving sum) on trials with no electrical stimulation and tested on four different sets of trials in which electrical microstimulation at four different times relative to saccade onset. E-D) Mean change in decoding performance for motor neurons (E) and visuomotor neurons (F) for each of the four stimulation condition. Each neuron’s performance was averaged over a window of −50 ms to −40 ms (“activation window”) from saccade onset. Error bars indicate s.e.m across neurons. *** indicates p < 0.001.

We calculated the difference of the time course of the decoders’ performance for stimulation and no-stimulation trials, separately for the four each microstimulation time periods tested (Fig. 4C-D). On visual inspection one sees fluctuations around zero for most of the conditions. The only exception is the performance for motor neurons. While also fluctuating, here the performance stayed below zero most of the time, indicating that stimulation compromised the saccade prediction.

In accordance with this visual judgment, the average of each neuron’s performance over the activation window revealed that the performance of motor neurons was significantly worse when the microstimulation was delivered shortly before saccade onset (Fig.4E; performance difference = −1.6%, p<0.001). On the other hand, the presaccadic code of visuomotor neurons did not show any sensitivity to the microstimulation.

## Discussion

In this study, we found that the activity of FEF motor neurons in a presaccadic window, supposedly representing the activation threshold for the initiation of saccade, was not dependent on the saccadic reaction times. A similar degree of independence was not exhibited by visuomotor neurons. Hence, motor neurons, yet not visuomotor neurons, satisfy the tenets of the variable rate model. We arrived at the same conclusion when resorting to a decoding analysis that avoided the many pitfalls of the correlation approach.

Several previous studies have suggested an independence of the discharge rate right before the saccade and reaction time based on zero correlations (Ding and Gold, 2012; Hanes and Schall, 1996; Jantz et al., 2013). One of them, (Jantz et al., 2013) suggested that there might be a difference between motor neurons and visuomotor neurons of the FEF, with only motor neurons complying with the variable rate model. In a simulation analysis, they showed that an ensemble of neurons with a strong sampling bias toward motor neurons would meet the predictions of the variable rate model. In fact, both our correlation and decoding analyses concur with their finding.

While manipulations of the input to FEF motor neurons will affect their discharge rates, they should not change the activation threshold for saccade initiation as it should reflect requirements of downstream neurons. In order to critically test this assumption, we carried out a microstimulation experiment in which the normal activity patterns of V4 neurons, providing visual information to the FEF was perturbated. In fact, the stimulation results did not match our expectations. Assuming that motor neurons would represent the FEF output, triggering the initiation of saccades by activating the premotor circuitry for saccades in the brainstem, their discharge before the initiation of saccades – in other words their activation threshold-should have been resilient to perturbation. Indeed, microstimulation delivered during the earlier periods before a saccade – more specifically, during baseline fixation, the presentation of the target and the subsequent delay period did not change activation thresholds of motor neurons. However, stimulation right before the saccade had a dramatic effect on the threshold, exhibited by the fact that the discharge rate in this late period was rendered unable to predict the saccade. As expected, the incapacity of the spiking of visuomotor to predict the initiation of a saccade, remained unaffected by the perturbation.

The existence of different types of neurons in FEF is well accepted based on the results of previous work (Lowe and Schall, 2018; Mitchell et al., 2007; Noudoost and Moore, 2011; Thiele et al., 2016). Within this framework it is also generally assumed that motor neurons are much closer to saccade initiation than visuomotor neurons (Hanes et al., 1998). On the other hand, to the best of our knowledge the relationship between the two different types of FEF neurons and other functions of the FEF such as the control of covert shifts of spatial attention (Moore and Fallah, 2004), visual stabilization (Chen et al., 2018), visual working memory (Merrikhi et al., 2017) or visual search (Yan and Zhou, 2019) is less clear, although the close relationship between these functions and the planning and execution of eye movements, i.e. overt shifts of attention, might suggest a similar architecture. Our correlation and decoding results indeed support a central role of motor neurons in saccade initiation according to the tenets of the variable rate model. However, at first glance the perturbation results seem to contradict notion that motor neurons control saccade initiation by crossing a preset and fixed activation threshold of recipient neurons in the brainstem saccade generator circuitry. However, the seeming contradiction resolves if we assume that screwing up the input to visuomotor neurons by stimulating V4 afferents may compromise the development of a normal saccade code at the level of motor neurons. In the same vein, also the activation thresholds of putative recipient neurons in the brainstem might be sensitive to changes of the normal spiking patterns, arguably induced by microstimulation of V4. In order to test these possibilities, experiments selectively manipulating visuomotor or motor neurons, e.g., by resorting to optogenetics, might provide pertinent information. Previous work has successfully identified brainstem projecting FEF neurons by retrogradely activating them from brainstem sites (Segraves and Goldberg, 1987). Recordings from the SC and eventually dependent sites in the lower brainstem might be needed to exclude a dependence of activation thresholds on the degree of naturalness of incoming spiking patterns.

## Acknowledgment

The authors would like to thank Dr. Behrad Noudoost for collecting the data used in this study and generously sharing it with us.

